## Supplementary notes and figures for "Self-organized intracellular twisters"

#### Materials and Methods

##### Simulation of microtubules in closed geometries

We briefly outline our method of simulation, the technical details of which appear in [1], which combine slender-body theory and boundary integral methods for solving the Stokes equations. A publicly available, and elaborated, version of the underlying code, *SkellySim*, is available at [2].

In broad strokes, we simultaneously solve the coupled equations of motion, Eqs. (1) and (2) in the main text, for the fluid and the immersed microtubules confined in the cellular volume $\Omega$ and clamped at the boundary $\Gamma$. Due to the linearity of the Stokes equations, we can write the fluid velocity at $\mathbf{x}$ in $\Omega$ as $\mathbf{u}(\mathbf{x}) = \mathbf{u}^{mt}(\mathbf{x}) + \mathbf{u}^{\Gamma}(\mathbf{x})$, with $\mathbf{u}^{mt}(\mathbf{x}) = \sum_i \mathbf{u}^i(\mathbf{x})$ the superposition of velocities induced by forces and conformations of each microtubule $i$, and $\mathbf{u}^{\Gamma}(\mathbf{x})$ the consequent backflow velocity induced by the no-slip condition at the confining boundary $\Gamma$. The velocities $\mathbf{u}^{mt}$ and $\mathbf{u}^{\Gamma}$ are expressed in terms of two fundamental solutions to the Stokes equations, the Stokeslet tensor $\mathbf{G}$ (a second-rank tensor), and the Stresslet $\mathcal{T}$ (a third-rank tensor):

$$\mathbf{G}(\mathbf{x}) = \frac{1}{8\pi\mu}\frac{\mathbf{I} + \hat{\mathbf{x}}\hat{\mathbf{x}}}{|\mathbf{x}|}; \quad \mathcal{T}(\mathbf{x}) = \frac{-3}{4\pi\mu}\frac{\hat{\mathbf{x}}\hat{\mathbf{x}}\hat{\mathbf{x}}}{|\mathbf{x}|^2}. \tag{S1}$$

$\hat{\mathbf{x}} = \mathbf{x}/\|\mathbf{x}\|$. Slender-body theory for the Stokes equations gives that, to leading (logarithmic) order in the slenderness ratio $\epsilon$, the velocity induced by a microtubule is given as a line integral of the distribution of the Stokeslets along its centerline:

$$\mathbf{u}^i(\mathbf{x}) = \int_0^{L^i} \mathbf{G}\left(\mathbf{x} - \mathbf{r}^i(s')\right)\mathbf{f}^i(s')\, ds'. \tag{S2}$$

where $\mathbf{f}_i$ is the internal elastic force that a microtubule exerts upon the fluid (see main text).

The second contribution, $\mathbf{u}^{\Gamma}$, accounts for the no-slip condition taken upon $\Gamma$, and is expressed as a surface convolution of the Stresslet over $\Gamma$ with an unknown density $\mathbf{q}$:

$$\mathbf{u}^{\Gamma}(\mathbf{x}) = \int_{\Gamma} dS_y\, \mathbf{n}(\mathbf{y}) \cdot \mathcal{T}(\mathbf{r}) \cdot \mathbf{q}(\mathbf{x}), \tag{S3}$$

where $\mathbf{r} = \mathbf{x} - \mathbf{y}$, and $\mathbf{n}$ is the outward normal vector to $\Gamma$. In the parlance of integral equations, this is a double-layer representation. In such a representation, taking the limit $\mathbf{x} \to \Gamma$ of Eq. (S1) and applying the no-slip condition, $\mathbf{u} = 0$, generates a well-conditioned Fredholm integral equation of the second kind for $\mathbf{q}$:

$$-\frac{1}{2}\mathbf{q}(\mathbf{x}) + \int_{\Gamma} dS_y \mathbf{n}(\mathbf{y}) \cdot \mathcal{T}(\mathbf{r}) \cdot \mathbf{q}(\mathbf{y}) + \int_{\Gamma} dS_y [\mathbf{n}(\mathbf{x})\mathbf{n}(\mathbf{y})] \cdot \mathbf{q}(\mathbf{y}) = -\mathbf{u}^{mt}(\mathbf{x}), \qquad \mathbf{x} \in \Gamma. \tag{S4}$$

Here, the last term of the RHS is added to complete the rank (i.e. make it uniquely invertible) of the integral equation. This term does not change the velocity $\mathbf{u}^{\Gamma}$ but does fix a constant in the pressure field [1]. In Eq. (1) of the main text the background velocity for microtubule $i$ is given by $\bar{\mathbf{u}}^i(\mathbf{x}) = \sum_{j \neq i} \mathbf{u}^j(\mathbf{x})$, i.e., the flows induced by all other microtubules. At each time, the unknown field to determine for microtubule $i$ is its tension field $T^i$ which enforces inextensibility. This condition generates, through Eq. (1) of the main text, integro-differential equations for all the $T^i$s. Solution of the coupled system for $(\mathbf{q}, T)$ allows calculation of the microtubule velocities $\mathbf{X}_t^i(s,t)$.

The integrodifferential operators along the centerlines of the microtubules are discretized using $4^{th}$-order finite differences. For the time stepping, we use an adaptive explicit/implicit backward time-stepping scheme, which maintains accuracy while removing high-order stability stiffness constraints from the bending term. This results in a dense linear system of equations which we solve using GMRES with block-diagonal preconditioners. We accelerate computing the hydrodynamic interactions using the Fast Multipole Method [3]. The complexity per time-step scales with the total number of discretization points, on microtubules and the cell surface.

#### Biophysical and numerical parameters of simulations

*Biophysical:* In our simulations, we chose the length of all microtubules to be $L = 20\ \mu$m – on the longer side if growing from dynamical instability – and having bending rigidity $E = 20$ pN $\mu$m$^2$ [4]. Given a microtubule diameter of $\sim 20$ nm gives $\epsilon \sim 5 \cdot 10^{-4}$. If the cell is spherical it is of radius $R = 100\ \mu$m, taken in this abstracted shape as the typical size for stage 10 *Drosophila* oocytes. For an "oocyte"-shaped cell, whose construction is described below, the length is 150 $\mu$m, and width 108 $\mu$m. The immersing fluid is taken as Newtonian with viscosity $\mu = 1$Pa s [5].

The relaxation time of a microtubule is estimated as $\tau_r = \eta L^4/E \sim 16,000$ s. For comparison, this is somewhat less than the duration of stage 10 of Drosophila development – approximately 10 hr=36,000 s – where large scale streaming flows first appear. We note that streaming persists into stage 12. Generally we have $(\tau_c, \tau_m) = (\tau_r/\bar{\rho}, \tau_r/\bar{\sigma})$. For beating case I, this gives $(\tau_c, \tau_m) \approx (3200, 180)$s, while for the streaming case II, we have $(\tau_c, \tau_m) \approx (1066, 355)$s.

*Numerical:* Microtubules are clamped orthogonally to the inner surface of a model cell. Microtubules are placed randomly on the cellular surface with a uniform probability, with their placements filtered to ensure that any two microtubules are not closer than distance $\Delta = 0.1L$ from each other. Each microtubule is discretized with 64 points. The maximum allowable time-step is $\Delta t = 0.16$ s, much smaller than any of faster time-scales $\tau_c$ or $\tau_m$.

#### Classification of microtubule dynamics in simulations

We established the phase diagram of the model based on the dynamics and shape of the microtubules in long-term simulations. In the stable phase, microtubules remain unperturbed and normal to the surface; in the beating phase, microtubules' shapes continuously change with time; in the streaming phase, microtubules attain steady deformed shapes. To classify the simulations into these three phases, we first measured the normalized positional variance of the microtubule's free end

$$\delta_i^2 = \frac{\left\langle (\mathbf{X}^i(L) - \langle \mathbf{X}^i(L) \rangle)^2 \right\rangle}{L^2}, \tag{S5}$$

where the averaging is over a period of $\Delta t = 10^{-3}\tau_r$. If $\delta_i^2 < 0.1$, we consider the microtubule shape time-independent; otherwise, its shape is dynamic. For a simulation, if more than 90% of microtubules are dynamic, we classify it as the beating phase; otherwise, it belongs to stable or streaming phases. To distinguish between these two phases, we measured the projection of the microtubule end-to-end vector as

$$a_i = 1 - \hat{\mathbf{n}} \cdot \frac{\mathbf{X}_i(L) - \mathbf{X}_i(0)}{|\mathbf{X}_i(L) - \mathbf{X}_i(0)|} \tag{S6}$$

If $a_i < 0.05$, the microtubule is considered normal to the surface, otherwise, it is deformed. For a simulation, if more than 90% of microtubules are normal to the surface, we classify it as the stable phase. Otherwise, we classify it as the streaming phase.

#### Analytical approximation of the streaming flow in sphere

We approximated the flow in simulations of the sphere as a superposition of a swirling flow, $\mathbf{u}_\mathrm{s}$, and an axisymmetric bitoroidal flow, $\mathbf{u}_\mathrm{t}$ [6]

$$\begin{aligned}
&\mathbf{u}_\mathrm{ana} = \mathbf{u}_\mathrm{s} + \mathbf{u}_\mathrm{t} \\
&\mathbf{u}_\mathrm{s}(r,\theta,\phi) = \Omega \frac{r}{R} \sin(\theta)\hat{\boldsymbol{\phi}} \\
&\mathbf{u}_\mathrm{t}(r,\theta,\phi) = \frac{A}{R^3}\left[r(r^2 - W^2)(1 - 3\cos^2\theta)\hat{\boldsymbol{r}} + r(5r^2 - W^2)\cos(\theta)\sin(\theta)\hat{\boldsymbol{\theta}}\right],
\end{aligned} \tag{S7}$$

where, $r$, $\theta$, $\phi$ are the radial, polar, and azimuthal coordinates in the sphere, and $\hat{\boldsymbol{r}}$, $\hat{\boldsymbol{\theta}}$, and $\hat{\boldsymbol{\phi}}$ are the unit vectors in the respective directions. The three parameters of the model are $\Omega$, the strength of the swirling flow, $A$, the strength of the bitoroidal flow, and $W$ is the radius associated with the bitoroidal flow. We fit our simulations of spherical geometry to this flow by minimizing $\xi = \int (\mathbf{u}_\mathrm{ana} - \mathbf{u}_\mathrm{sim})^2 dV$, where the integration is over the sphere volume. For the minimization, we use the gradient descent algorithm for six free parameters, including three angles, to align the axis of the flow in simulation to the $z$ axis. We found that $\Omega = (100.2 \pm 3.0)$ nm/s, and $A = (5.5 \pm 1.2)$ nm/s. The ratio of the strength of the swirling flow to the toroidal one is $\Omega/A \sim 20$.

#### Simulation in oocyte-shaped geometry

To study the model in a geometry similar to *Drosophila* oocyte, we construct a surface of revolution as

$$\begin{aligned}
X &= Dx \\
Y &= Dr\cos(\phi) \\
Z &= Dr\sin(\phi),
\end{aligned} \tag{S8}$$

where $x \in (0,1)$, $\phi \in [-\pi,\pi)$, and $r = \frac{Tx^{p_1}(1-x)^{p_2}}{2^{(1-p_1-p_2)}}$(see [7]). $l$ is the oocyte length, $T$ sets the aspect ratio of the oocyte, and the parameters $p_1 \in [0,1]$ and $p_2 \in [0,1]$ determine the local curvature of the oocyte. In our simulations, we chose $D = 150\mu m$, $T = 0.72$, $p_1 = 0.4$, and $p_2 = 0.2$.

#### Live imaging of the *Drosophila* oocyte

Young mated female adults were fed with dry active yeast for 16-18 hours and dissected in Halocarbon oil 700 (Sigma-Aldrich, Cat: H8898) as previously described [8, 9]. Samples were imaged within 1 hour after dissection, using Nikon W1 spinning disk confocal microscope (Yokogawa CSU with pinhole size 50 µm) with Photometrics Prime 95B sCMOS Camera or Hamamatsu ORCA-Fusion Digital CMOS Camera, and a 40X 1.25 N.A. silicone oil lens, controlled by Nikon Elements software. 3D time-lapses were acquired every 10 seconds at $1\mu m$/step.

Flies were maintained on standard cornmeal food (Nutri-Fly Bloomington Formulation, Genesee, Cat: 66-121) supplemented with dry active yeast (Red Star) at room temperature (24 – 25°C). The following fly stocks were used in this study: mat $\alpha$tub-Gal4[V37] (III, Bloomington *Drosophila* Stock Center:7063) [10]; UASp-F-Tractin-tdTomato (II, Bloomington stock center:58989) [10, 11]; GFP::$\alpha$tub [12].

#### Reconstruction of 3D velocity field from live imaging

Here, we describe the steps for 3D reconstruction of the velocity field and measurement of microtubule orientation from experimental images. First, we reconstructed the 3D oocyte periphery, then measured the 2D cytoplasmic velocity field for each z-plane using particle image velocimetry, and then used it to reconstruct the 3D velocity field. We also measured the local orientation of microtubules using a linear filter for texture analysis.

#### 3D Reconstruction of the oocyte periphery

We developed an active contour method [13,14] to partially reconstruct the 3D geometry of the oocyte from volumetric images of F-actin. We first segmented the oocyte periphery for each z-plane and then used these to reconstruct the 3D oocyte surface. In short, for the middle z plane, we provided a closed curve, $\tilde{\Gamma}(s)$, which serves as the initial guess for the active contour method. The shape of the oocyte in this z-plane is given by minimizing the cost function

$$E[\Gamma(s)] = \oint_{\Gamma(s)} \frac{\alpha}{2}|\Gamma_{ss}|^2 ds + \iint_R (I_N(x,y) - \beta)dxdy \tag{S9}$$

where $\Gamma_{ss} = \partial^2\Gamma/\partial s^2$. The first term accounts for the smoothness of the contour, and the second term accounts for the interaction of the contour with the image, where $\beta$ is set such that the contour expands if it is far from the periphery. The image intensity, $I_N(x,y)$, is the normalized smoothed gradient of the F-actin image. We then used the segmented shape of the oocyte in this z-plane as an initial guess to segment the oocyte periphery in consecutive z-planes.

#### 2D particle image velocimetry

Particle image velocimetry (PIV) is a common technique for inferring the local velocity of the fluid by measuring the displacement of tracer particles between two consecutive time points. In brightfield microscopy images of *Drosophila* oocyte, lipid granule particles have high contrast relative to the cytoplasm and can serve as tracer particles to measure local cytoplasmic velocity. We developed a platform to perform PIV on brightfield microscopy images of *Drosophila* oocytes. A key piece of our software is using the contrast-limited adaptive histogram equalization method to enhance the contrast of the brightfield images [15]. To accurately measure the velocity in the complex geometry of the oocyte, we combined fast Fourier transform-based PIV (FFT-based) on a square grid within the interior of the oocyte and correlation-based PIV near the periphery.

For FFT-based PIV, square boxes of 100 pixels with 20 pixels spacing far from the oocyte periphery were taken [16,17]. For each box, we calculated the Fourier transform of the image intensity for two constitutive time points, $\tilde{I}_t(u,v)$ and $\tilde{I}_{t+1}(u,v)$, calculated the Hadamard product of one with the complex conjugate of the other, $\tilde{I}_H = \tilde{I}_t \circ \tilde{I}^*_{t+1}$, and set the displacement in that box as the position of the maximum of the inverse Fourier transform of $I_H(\Delta x, \Delta y)$ (Fig. S2a blue; pixel size, $0.260\mu m$).

For the correlation-based PIV for points near the periphery, we first constructed grids with shapes derived from the oocyte outline as follows: we chose $N$ evenly spaced points on the periphery, and for each point, we constructed $M$ evenly spaced points on a line connecting the center of mass of the oocyte cross-section to that point. By connecting each set of points, we constructed $N \times M$ grids (Fig. S2a). For each grid, we calculated the displacement by finding the maximum of the correlation function

$$H(\Delta_x, \Delta_y) = \int\int_A I_t(x,y) I_{t+1}(x + \Delta_x, y + \Delta_y)\mathrm{d}x\mathrm{d}y, \tag{S10}$$

where $I_t$, and $I_{t+1}$, are the mean subtracted intensity, and the integration is over the area of the grid. If there are multiple local maxima, we choose the one giving a smooth displacement field between the neighboring grids. Finally, we use interpolation to estimate the planar components of the velocity field, $u_x(x,y,z)$ and $u_y(x,y,z)$, on a regular grid across the oocyte using these two displacement fields (Fig. S2a).

#### Approximation of the out-of-plane velocity

We measured the out-of-plane component of the velocity field, $u_z(x,y,z)$, by assuming the incompressibility of the cytoplasm and the impermeability of the oocyte boundary ($\Gamma$) (on the timescales of microscopy) and solving

$$\nabla \cdot \mathbf{u} = 0; \quad \mathbf{u} \cdot \hat{\mathbf{n}}|_\Gamma = 0 \tag{S11}$$

where $\hat{\mathbf{n}}$ is the surface normal vector. To do so, we numerically solve the ODE

$$\frac{\partial u_z}{\partial z} = -\left(\frac{\partial u_x}{\partial x} + \frac{\partial u_y}{\partial y}\right), \tag{S12}$$

with the boundary condition $u_z = -(n_x u_x + n_y u_y)/n_z$ at the oocyte periphery (Fig. S2b right).

#### Estimation of microtubule orientation field from microscopic images

To measure the local microtubule orientation, we use a Gabor filter, which is a linear filter for texture analysis [18]. It allows examination of any specific frequency content in the image in a given direction. The inputs of the filter are its wavelength and orientation, and the outputs are the magnitude and phase response to the filter. We use 3-pixel wide wavelength for angles $\theta \in (0, \pi]$ with interval $\pi/180$, and for each angle, we calculate the magnitude response in grids of $20 \times 20$ pixels. We set the grid orientation as the angle with the largest magnitude response and grid magnitude as the value of the magnitude response to that angle (Fig. S4).

#### Estimation of the positions of the defect

As illustrated in Fig. 2E in the main text, near the defect centers microtubules are relatively straight and normal to the surface. To find the defect positions in the oocyte simulation, we sort all the microtubules in the descending order of $a_i$ (equation (S6)), the length of projection of the end-end vector on the surface. We assign the first defect center to be at the location where the microtubule with lowest $a_i$ is clamped. From the rest of the microtubules, we find the one with next lowest $a_i$, which is at least 40 $\mu m$ away from the first defect and assign it's clamping position to be the second defect.

### Supplementary movie captions

Movie 1: Time-course of configurations of microtubules anchored to the interior surface of a sphere in a cut-away view for a simulation with parameters $\bar{\rho}$=5 and $\bar{\sigma}$=90 (Case I). Timestamp shows time normalized to the relaxation time of a single microtubule.

Movie 2: Time-course of 2D projection of velocity field for a simulation with parameters $\bar{\rho}$=5 and $\bar{\sigma}$=90 (Case I). The bordering circle represents the boundary of the sphere (Case I). Timestamp shows time normalized to the relaxation time of a single microtubule.

Movie 3: Time-course of configurations of microtubules anchored to the interior surface of a sphere in a cut-away view for a simulation with parameters $\bar{\rho}$=15 and $\bar{\sigma}$=45 (Case II). Timestamp shows time normalized to the relaxation time of a single microtubule.

Movie 4: Time-course of 2D projection of velocity field for a simulation with parameters $\bar{\rho}$=15 and $\bar{\sigma}$=45 (Case II). The bordering circle represents the boundary of the sphere (Case II). Timestamp shows time normalized to the relaxation time of a single microtubule.

Movie 5: Live movie of a cross-section of *Drosphila* oocyte obtained by a brightfield microscope. The scale bar denotes a length of 50 $\mu m$. The timestamp is in unit of minutes.

Movie 6: Time-course of the polar order parameter $P$ for a simulation with parameters $\bar{\rho}$=15 and $\bar{\sigma}$=45. Left and right views face two diametrically opposite defect like structures formed in long time. Timestamp shows time normalized to the relaxation time of a single microtubule.

Movie 7: $360^0$ rotation of two snapshots at early time ($t = 0.025\tau_r$, $t = 0.035\tau_r$) and one snapshots at long time ($t = 0.5\tau_r$) of a simulation with parameters $\bar{\rho}$=15 and $\bar{\sigma}$=45. $\tau_r$ is the relaxation time of a single microtubule. Surface polarity vectors $p_i$ corresponding to each microtubule are represented as arrows and superposed on the surface. The color of the surface represents the polar order parameter $P$.

Movie 8: Time-course of configurations of microtubules anchored to the interior surface for a cell shape similar to that of an oocyte, shown in two cut-away views. The simulation parameters are $\bar{\rho}$=15 and $\bar{\sigma}$=45. Timestamp shows time normalized to the relaxation time of a single microtubule.

### Supplementary figures

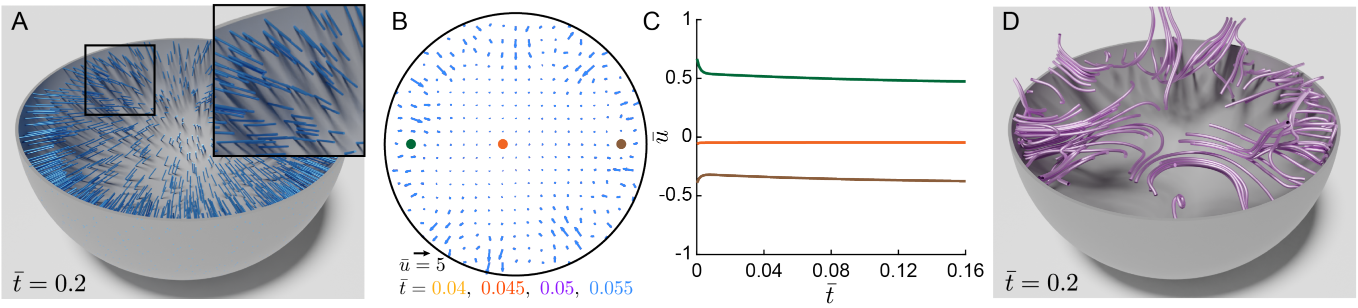


Figure S1: Simulation in the stable regime, using $\bar{\rho} = 15$ and $\bar{\sigma} = 5$. (A) Cut-away view of instantaneous microtubule configurations in the spherical cell. Inset shows very slightly bent microtubules. (B) 2D projection of velocity field in the sectioning equatorial plane of (A), at four time points. (C) the azimuthal velocity component at the three points labelled in the equatorial plane, as a function of $\bar{t}$. Note the very small magnitude of the velocities. (D) 3D streamlines integrated from the (low magnitude) 3D velocity field for a simulation for a stable regime with parameters .

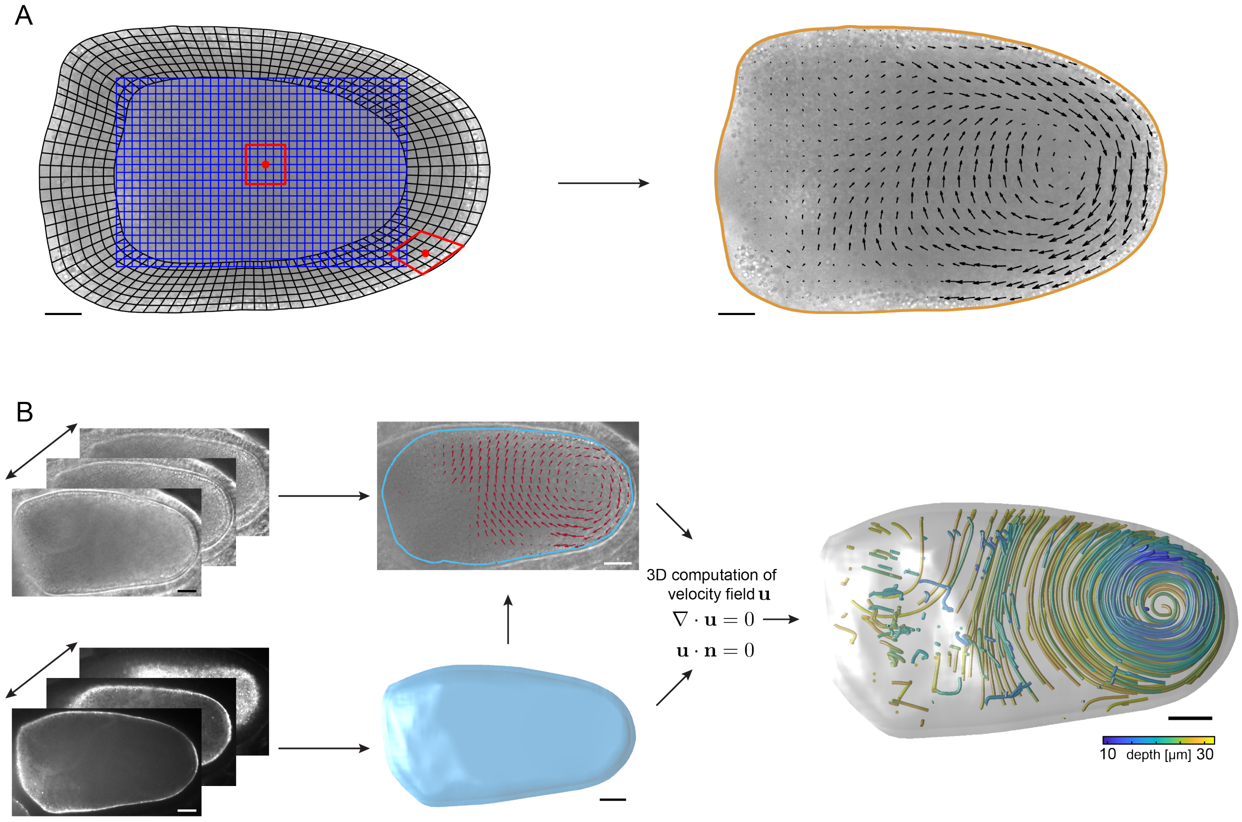


Figure S2: Elements of the 3D flow measurements in the *Drosophila* oocyte. (A) Left panel shows the overset grid generated by combining a Cartesian mesh for points far from oocyte boundary (blue) and a boundary-conforming mesh constructed from segmentation of the oocyte periphery (black). Right panel shows the 2D cytoplasmic velocity field in the corresponding slice. Scale bar, $25\mu m$. (B) Steps for the reconstruction of 3D flow structure from live images: 3D segmentation of oocyte periphery, 2D PIV on individual slices, computation of out-of-plane velocity component by assuming flow incompressibility, and 3D flow streamline reconstruction. Images are as seen in the microscope, with anterior at the left and posterior at the right. Scale bar, $25\mu m$.

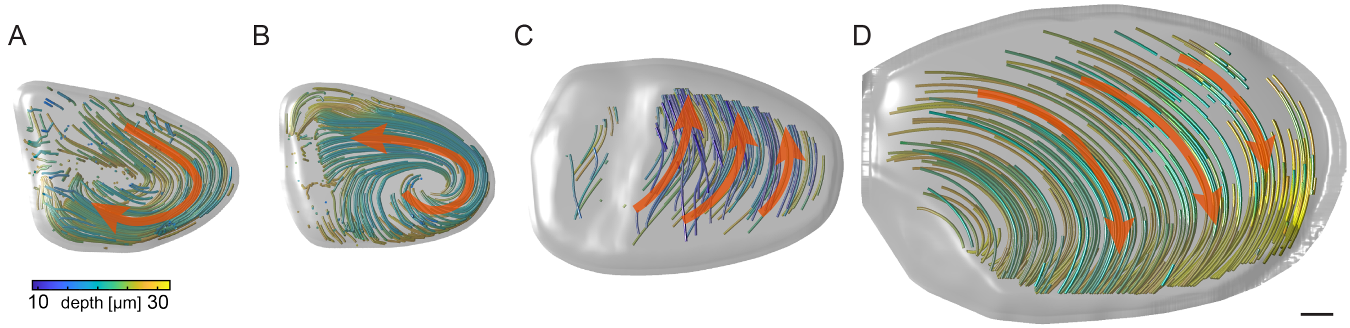


Figure S3: Examples of flow structures observed within different experimental oocytes. (A,B) show vortical "twister-like" flows, while (C,D) show likewise commonly observed cross-cell streaming. Red arrows show flow direction. All images are as seen through the microscope, and are oriented with anterior at the left, and posterior at the right. Note that if interpreted through the lens of a confined twister within the cell, (C) could be interpreted as counter-clockwise motion around the oriented anterior-posterior axis, while (D) takes the opposite direction. Scale bar, $25\mu m$.

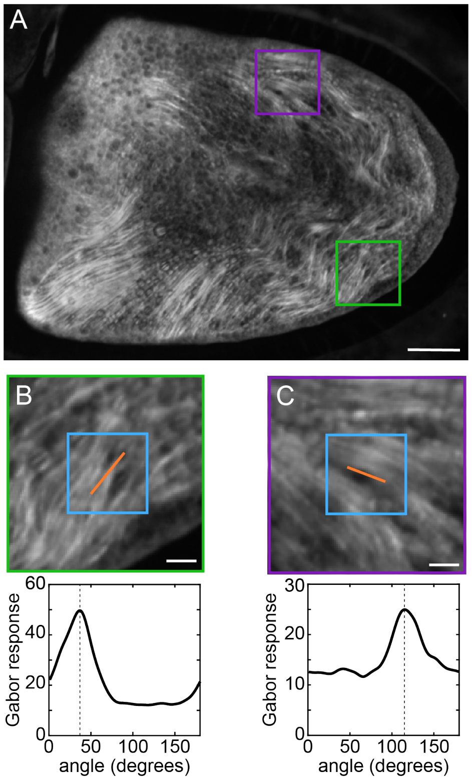


Figure S4: Measurement of microtubule orientation from live images of *Drosophila* oocytes. (A) A 2D projection of microtubules tagged with maternally derived GFP-$\alpha$tub. Scale bar, $25\mu m$. (B)-(C) B corresponds to the green box in A, while C corresponds to the purple. Orange lines: microtubule orientation of peak power response of the Gabor filter applied to a $20 \times 20$ pixel box (blue). The graphs are the Gabor response as a function of angle relative to the x-axis. Images are as seen in the microscope, with anterior at the left and posterior at the right. Scale bar, $5\mu m$.

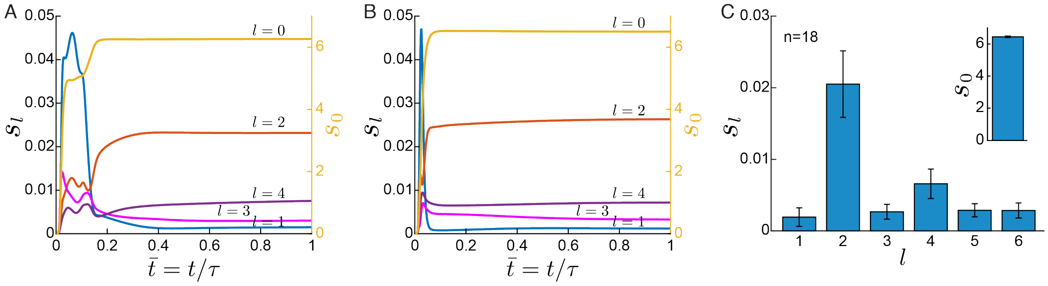


Figure S5: Robustness of twister formation to microtubule distribution. (A-B) Angular power $s_l$ of spherical harmonic coefficients of the $P$ field, $s_l(t) = \sum_{m=-l}^{m=l} |\tilde{P}_{lm}(t)|^2/(2l+1)$, for two simulations with different microtubule anchoring distributions but with same (straight) initial configuration. The right and left axes represent $l = 0$ and $l \neq 0$, respectively. (C) Bar graphs representing the mean steady state value of $s_l$ ($l = 0$ in insets) for 18 such simulations (error bars show standdard deviations).

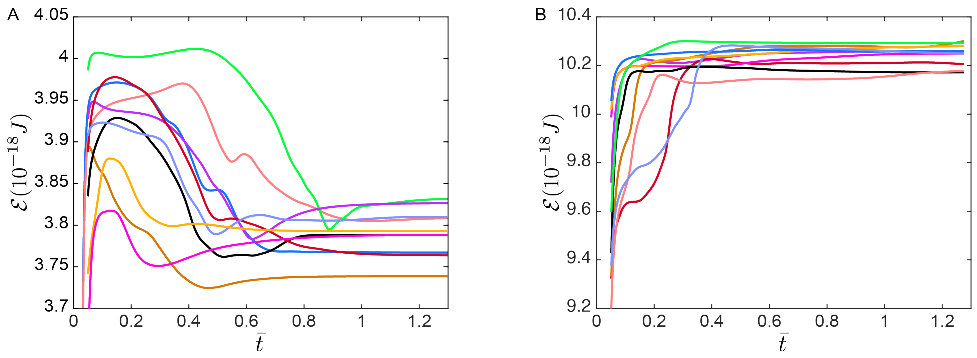


Figure S6: Evolution of the elastic energy in spherical and oocyte-shaped cells. (A) total elastic energy $\mathcal{E}$ for 10 independent realizations (different microtubule placements) of case II in a spherical cell geometry as a function of adimensional time $\bar{t} = t/\tau_r$. Note that all simulations evolve towards twisters of similar elastic energy. (B) 10 such simulations, again using case II parameters, in the oocyte geometry of Fig. 4E of the main text. All show an overshoot in the elastic energy, followed by relaxation to a lower energy axisymmetric state. These simulations correspond to those shown in Fig. 4F of the main text demonstrating reorientation to axisymmetry.
